# An almost chromosome-level assembly and annotation of the *Alectoris rufa* genome

**DOI:** 10.1101/2024.01.11.575009

**Authors:** Abderrahmane Eleiwa, Jesus Nadal, Ester Vilaprinyo, Alberto Marin-Sanguino, Albert Sorribas, Oriol Basallo, Abel Lucido, Cristobal Richart, Romi Pena, Roger Ros-Freixedes, Anabel Usie, Rui Alves

**Affiliations:** Institut de Recerca Biomédica (IRBLleida); Universitat de Lleida (UdL); Universitat Rovira i Virgili (URV); Centro de Biotecnologia Agrícola e Agro-alimentar do Alentejo (CEBAL)/Instituto Politécnico de Beja (IPBeja); MED–Instituto Mediterrâneo para a Agricultura, Ambiente e Desenvolvimento & CHANGE–Global Change and Sustainability Institute

## Abstract

The red-legged partridge, *Alectoris rufa* (n=38 chromosomes) plays a crucial role in the ecosystem of southwestern Europe, and understanding its genetics is vital for conservation and management. Here we sequence, assemble, and annotate a highly contiguous and nearly complete version of it genome (115 scaffolds, L90=23). This assembly contains 96.9% (8078 out of 8332) orthologous genes from the BUSCO aves_odb10 dataset of single copy orthologous genes. We identify RNA and protein genes, 95% of which with functional annotation. This near-chromosome level assembly revealed significant chromosome rearrangements compared to quail (*Coturnix japonica*) and chicken (*Gallus gallus*), suggesting that *A. rufa* and *C. japonica* diverged 21 M-years ago and that their common ancestor diverged from *G. gallus* 37 M-years ago. The reported assembly is a significant step towards a complete reference genome for *A. rufa*, contributing to facilitate comparative avian genomics, and providing a valuable resource for future research and conservation efforts for the red-legged partridge.

## INTRODUCTION

*Alectoris rufa*, also known as red-legged partridge, is a game bird that holds significant ecological and economic importance for rural areas in southwestern Europe ^1^. Habitat degradation, captive breeding, and hunting management have led to the creation of a complex species situation, impacting both the ecosystems and society of the region. Across various hunting grounds, wild, farmed, and hybrid partridges coexist in varying proportions. While these partridges exhibit distinctions in behavior, physiology, morphology, anatomy, and genetics, the absence of a reference genome hinders our ability to molecularly differentiate these ecotypes, spanning from wild to domestic ^2^. The haploid genome of *A. rufa* has 10 macro chromosomes, two of which are sex chromosomes, and 30 micro chromosomes ^3,4^. The advent of Next-Generation Sequencing (NGS) technologies, based on short-read sequencing data, combined with decreasing DNA sequencing costs, led to an increase of the number of available genome sequences. However, those genomes were still highly fragmented due to the limitations inherent to short reads, where for example repetitive regions can lead to genome misassembles. The emergence of third-generation sequencing technologies partially overcame those limitations by generating long-read sequencing data. These long-reads helped reduce assembly fragmentation and increase contiguity, greatly improving the quality of whole-genome assemblies ^5^. Still, early long-read technologies have base-calling error rates of 10-14%, that are much higher than the less than 1% error rate found in short-read technologies ^6^. In addition, error profiles in both technologies are different. Errors in short-reads are mostly at the level of wrong nucleotide substitutions, while errors in long-reads mostly involved incorrect insertions and deletions ^7,8^. This difference makes long read errors more complex to resolve, requiring an error correction step prior to genome assembly. The error correction problem has been addressed either by self-correction, aligning the long-reads against each other, or by a hybrid approach in which the long-reads are corrected using short-reads. The later approach is known to better achieve accurate genome assemblies when compared against genomes assembled based only on short-or long-read technologies ^9,10^.

In this context, the quality of reference genome assemblies benefited from the combination of Illumina short-read sequencing with third-generation sequencing platforms such as Pacific Bioscience (PacBio) ^11^ or Oxford Nanopore Technologies (ONT) ^12^. Application of these technologies improved contiguity, completeness, and accuracy compared to assemblies based on short-read sequencing alone ^13,14^. In general, the number of contigs and scaffolds was significantly reduced, and N50 values increased, leading to better genome annotation and identification of more genes, including non-coding RNA genes, pseudogenes, and transposable elements ^15,16^. Examples of genomes assembled using hybrid approaches in the avian clades include, for example, tibetan partridge ^15^, indian peafowl ^17^, domestic turkey ^14^, and the common pheasant ^18^.

The first effort to sequence the red-legged partridge genome of a male individual, which was published in 2021 under the accession number GCA_019345075.1 ^19^, was based on Illumina paired-end short reads sequence data resulting in a highly fragmented assembly, with 10 598 scaffolds, a contig/scaffold N50 of 11.57 Mb, and L90 equal to 131. A more recent version of *A. rufa*‘s genome, based on ONT reads, was recently released at the NCBI under the accession number of GCA_947331505.1. That version has 426 scaffolds with N50 of 34 Mb and L90 of 32 (unpublished work). Both genomes lack detailed annotation, and significant variations exist between the two assemblies. These discrepancies may lead to potential errors in gene order conservation (synteny) and contribute to large-scale assembly inaccuracies. In order to overcome some of the challenges and limitations found in the earlier genome assemblies of *A.rufa* and move towards a well annotated chromosome level assembly, we combine short- and long-read sequencing data in a hybrid approach. Here we report the resulting almost chromosome level assembly and provide a high-quality genome annotation. We validated the assembly by comparing it to the reference genomes of chicken (*Gallus gallus*, NCBI reference GCF_016699485.2) and quail (*Coturnix japonica,* NCBI reference GCF_001577835.2), two closely related species. Overall, we provide a valuable resource for comparative and population genomics, improving our understanding of avian evolution, biogeography, and demography.

## RESULTS

### Estimation of genome size and heterozygous rate

We conducted genome profiling on sixty *A. rufa* individuals using *k*-mer analysis of short-read sequence data and finding an estimated genome size between 1 and 1.06 Gb, and 0.1%≤heterozygosity≤0.4% (Figure 1A).

**Figure 1:**
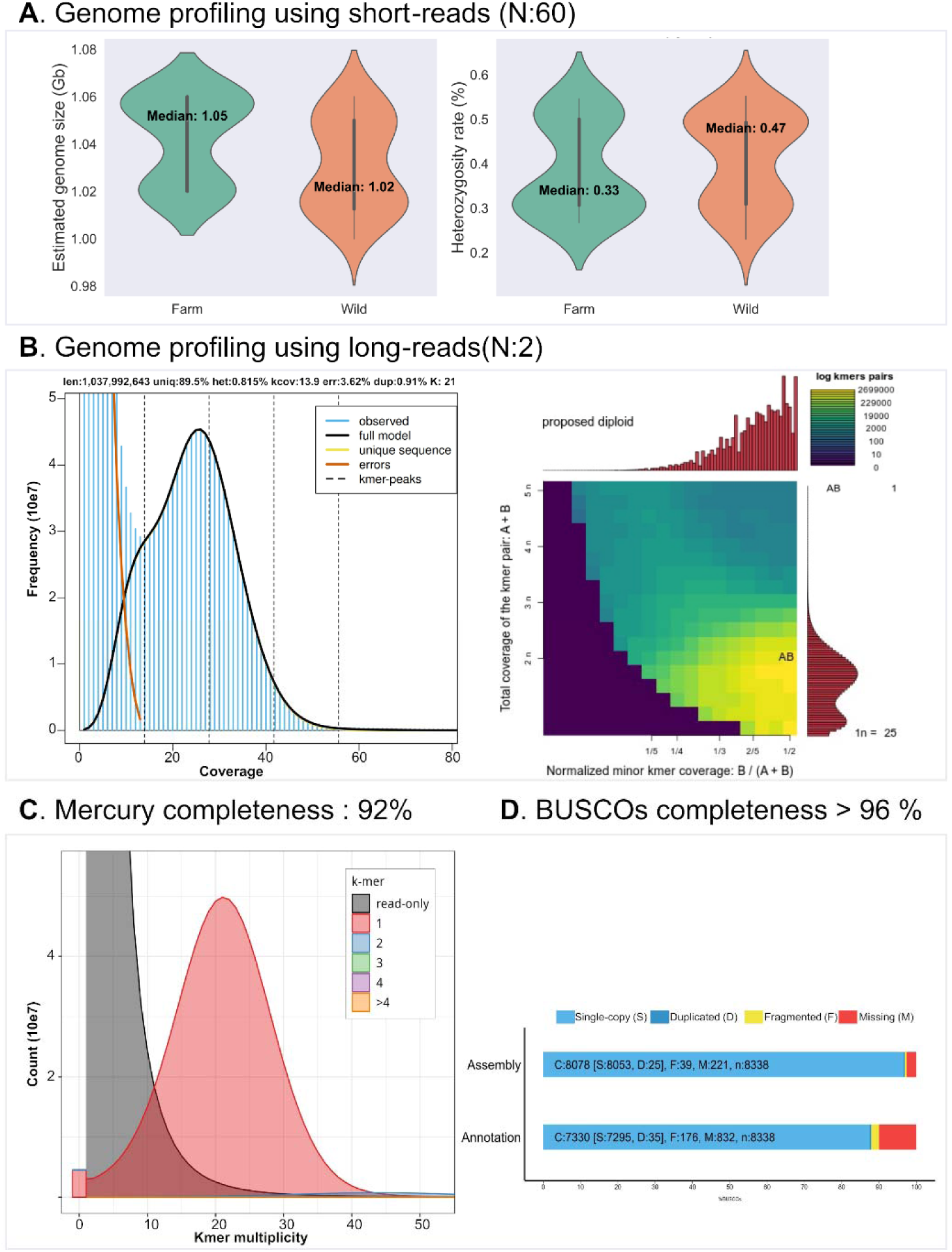
Assessment of *de-novo* assembly and genome profiling in *Alectoris rufa*. **A –** Distribution of genome size and heterozygosity levels estimated by jellyfish and Genomescope2 across sixty individuals of *A. rufa* sequenced with Illumina short-reads, including 30 domesticated (farm) and 30 wild specimens. B **–** Genome size and ploidy level determination using long reads from two individual *A. rufa* sequenced with ONT. Left: representation of genome size and heterozygosity using GenomeScope2 plot. Right: genome ploidy inference using Smudgeplot approach. C **–** Assessment of scaffolds completeness through the k-mer spectrum plot approach by Mercury, indicating scaffold completeness based on a k-mer size of 21 were 92%. D **–** Evaluation of assembled scaffold completeness based on recovered core genes from the aves_odb10 dataset of BUSCO.

### *A. rufa* genome assembly, annotation and quality assessment

We tested and evaluated various pipelines to assemble the genome of the red-legged partridge. The NextDenovo pipeline produced a primary assembly with the best metrics. This assembly comprised 116 contigs, with an N50 length of 74 and an N90 of 10 Mb (Table S1). We further refined this assembly, increasing the number of recovered single-copy genes found in the BUSCO dataset of avian single copy orthologous genes (aves_odb10, N= 8332 genes) from 90.6% (7555 out of 8338) to 96.8% (8078 out of 8332) (Table S2). The contigs were then used as the basis for genome scaffolding, resulting in a final genome assembly of 115 scaffolds and 1.03 Gb. Table 1 summarizes the most relevant contiguity metrics of this assembly and its annotation.

**Table 1:** Statistic of the red legged partridge (*Alectoris rufa*) genome de novo assembly and annotation.

|  | *Alectoris rufa* |
| --- | --- |
| **Genome assembly** |  |
| Assembly size (Gb) | 1.03 |
| Number of scaffolds | 115 |
| Max. scaffold length (Mb) | 149.6 |
| N50 scaffold size (Mb) | 74 |
| N90 | 23 |
| Mean scaffold length (Mb) | 9.03 |
| QV | 36.3 |
| Mercury completeness score |  |
| **Gene annotation** |  |
| Total Protein-coding genes | 30236 |
| **Functional annotation (Total)** |  |
| Uniprot (SWISS_prot) | 18878 |
| Uniprot (TrEMBL) | 28596 |
| NCBI_NR | 28799 |
| **Genomic features** |  |
| Repeat (%) | 13.3 |
| GC (%) | 42.1 |
| Coding (%) | 47.3 |
| Intron (%) | 43.4 |
| Gene length (bp) | 485558167 |
| Avg. protein length (aa) | 441.3 |
| Total microRNAs | 246 |
| Total tRNA | 305 |
| Total rRNA | 130 |
| Total snRNA | 315 |

The final assembly significantly improves the statistical metrics of contiguity of the earlier available assemblies (Table S2). Our L90 is 23, close to the 10 macro-chromosomes present in the haploid genome of *A. rufa*, and at least five times smaller than that for assemblies GCA_947331505.1 (based on short-reads) and GCA_019345075.1 (based on long-reads). Our N50 (74 Mb) is twice that of the GCA_019345075.1 assembly and seven times that of the GCA_947331505.1 assembly. Our assembly contained 96.78% (n=8053 genes) of complete and single-copy genes without duplications present in the BUSCO avian dataset, surpassing both the short-read (95.1%; n=7933 genes) and the long read (96.58%; n=7378) genome assemblies. Table 2 summarizes the main difference in terms of the genome contiguity and completeness metrics between those assemblies.

**Table 2:** Sequencing information and the quality metric of our assembly compared with other previously published assembly of the *Alectoris rufa* genome.

|  | **This study** | **Chattopadhyay et al, 2021** | **GCA_947331505.1** |
| --- | --- | --- | --- |
| **Sequencing technology** | Illumina<br>NovaSeq 6000,<br>ONT GridION | Illumina<br>HiSeqX | ONT GridION |
| **Assembly size (bp)** | 1027480606 | 1039068021 | 1142486555 |
| **Total scaffolds** | 115 | 10598 | 426 |
| **Scaffold N50 (bp)** | 74759052 | 11577318 | 37566138 |
| **Scaffold N90 (bp)** | 10103736 | 1025037 | 6380696 |
| **Longest Scaffold length (Mb)** | 149 | 47 | 118 |
| **GC content (%)** | 42.1 | 41.4 | 42.3 |
| **Total predicted protein-coding genes** | 30236 | NA | NA |
| **Complete recovered BUSCOs** | 8078 | 7970 | 8094 |
| **Complete and single-copy BUSCOs** | 8053 | 7933 | 7378 |
| **Complete and duplicated BUSCOs** | 25 | 37 | 716 |
| **Fragmented BUSCOs** | 39 | 122 | 37 |
| **Missing BUSCOs** | 221 | 246 | 207 |
| **Total BUSCO groups searched** | 8338 | 8338 | 8338 |

ONT = Nanopore Oxford Technology.

Additionally, we compared our genome assembly against that of eleven birds and one reptile, all of which possessed chromosome-level genome assemblies (Table S3). Our assembly has the third highest N50 value, surpassed only by the *G. gallus (chicken)* and *Anas platyrhynchos (mallard)* genomes (Figure 2A). Moreover, in terms of avian orthologs, our assembly ranks within the top five of the highest number of both complete and single copy orthologs. As expected, the genome of the reptile *Anolis carolinensis*, used subsequently as outgroup for phylogenetic comparisons, exhibited the lowest presence of avian orthologs (Figure 2B). The NCBI’s Foreign Contamination Screening revealed no significant contamination in the assembly of those 115 scaffolds (Table S4).

**Figure 2:**
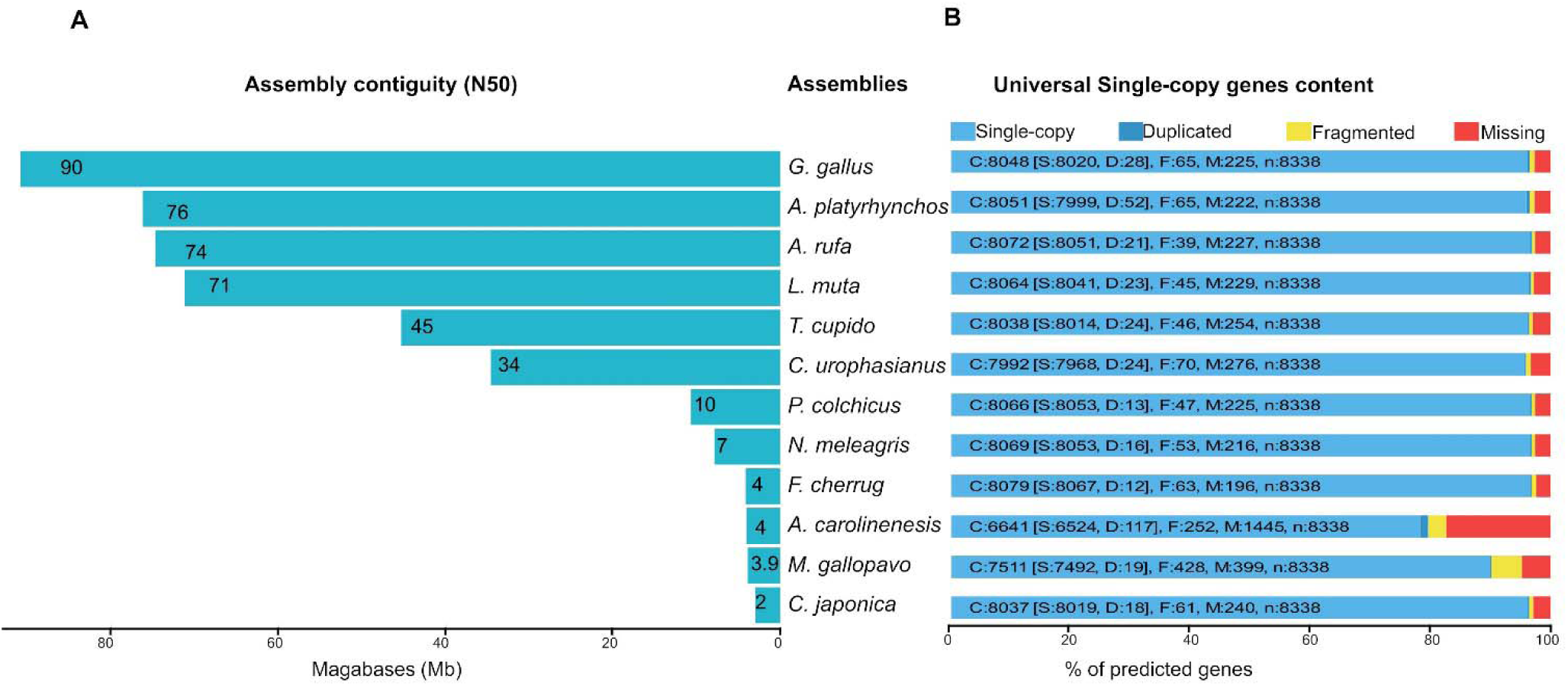
A comprehensive assessment of the completeness and correctness at the chromosome level of the *A. rufa* assembly in comparison to closely related bird species and using the reptile *A. carolinensis* as outgroup. A – N50 statistic for each genome assembly. B – Completeness of each assembly based on BUSCO results with the aves_odb10 dataset.

### Annotation of transposable elements

RepeatMasker ^20^ annotated 13% of the *A. rufa* genome as repetitive sequences. Table 3 summarizes the analysis of transposable elements (TE), which revealed a higher percentage of repetitive elements, when compared to the previous draft genome based on short reads alone ^19^. Long interspersed nuclear elements (LINE) are the most frequent transposable elements in the genome, representing 7.74% of the whole genome sequence. DNA transposons (2.33%) and long terminal repeat (LTR) elements (1.76%) are the second and third most abundant classes of transposable elements in the genome.

**Table 3.** Statistics of repetitive element in the current assembly compared to previous short reads alone-based assembly.

|  | **This study** |  | **Chattopadhyay et al, 2021** |  |
| --- | --- | --- | --- | --- |
|  | **Length (bp)** | **% of genome** | **Length occupied** | **% of genome** |
| SINEs | 723913 | 0.07 | 179454 | 0.02 |
| LINEs | 80386206 | 7.74 | 70223435 | 6.83 |
| LTR elements | 18335204 | 1.76 | 10566378 | 1.03 |
| DNA transposon | 24247058 | 2.33 | 7824132 | 0.76 |
| Unclassified | 14562721 | 1.40 | 10691987 | 1.04 |
| Total | 138255102 | 13.31 | 99485386 | 9.68 |

SINE: short-interspersed element; LINE: long-interspersed elements; LTR: long terminal repeat.

### Annotation of RNA and protein-coding genes

We annotated 30236 protein-coding genes in our assembly, spanning ∼46% of the genome (Table 4). The average length of annotated protein-coding gene was 16058 bp, with an average exon size of 194 bp and intron size of 2533 bp (Table 4). Out of the total predicted 30236 gene models, 8509 genes were validated using transcriptomic data from spleen samples (Table S5).

**Table 4:** Statistics summary of genomic features in term of genomic length structurally annotated of the *A. rufa* genome.

| **Genomics features** | **Counts** |
| --- | --- |
| Number of genes | 30236 |
| Number of CDS | 30236 |
| Number of exons | 206051 |
| Number of intron in CDS | 175815 |
| Number gene overlapping | 0 |
| Number of single exon gene | 8625 |
| mean mRNAs per gene | 10 |
| mean exons per CDS | 68 |
| mean introns in CDSs per mRNA | 58 |
| Total gene length (bp) | 485558167 |
| Total CDS length (bp) | 40122173 |
| Total intron length per CDS | 445435994 |
| mean gene length (bp) | 16058 |
| mean CDS length (bp) | 1326 |
| mean CDS piece length (bp) | 194 |
| mean intron in CDS length (bp) | 2533 |

By leveraging eggNOG-based functional annotation, we identified homologs for 83.9% (23398) of the predicted protein genes. 57.1% (13371) of the predicted protein genes were assigned specific biological functions based on the gene ontology (GO) classification (Supplementary data file S1). Then, we blasted the full gene set to the SwissProt, TrEMBL and NCBI protein databases. Remarkably, 95% (28862) of the predicted protein-coding genes significantly matched homologous sequences in at least one of the three databases. Of these, 18865 (62.1%) proteins were simultaneous and consistently annotated between the three databases (Figure 3). Additionally, employing InterProScan, we successfully assigned 25978 (85.9%) predicted protein-coding genes to specific family and subfamily domains (Supplementary data file S1).

**Figure 3:**
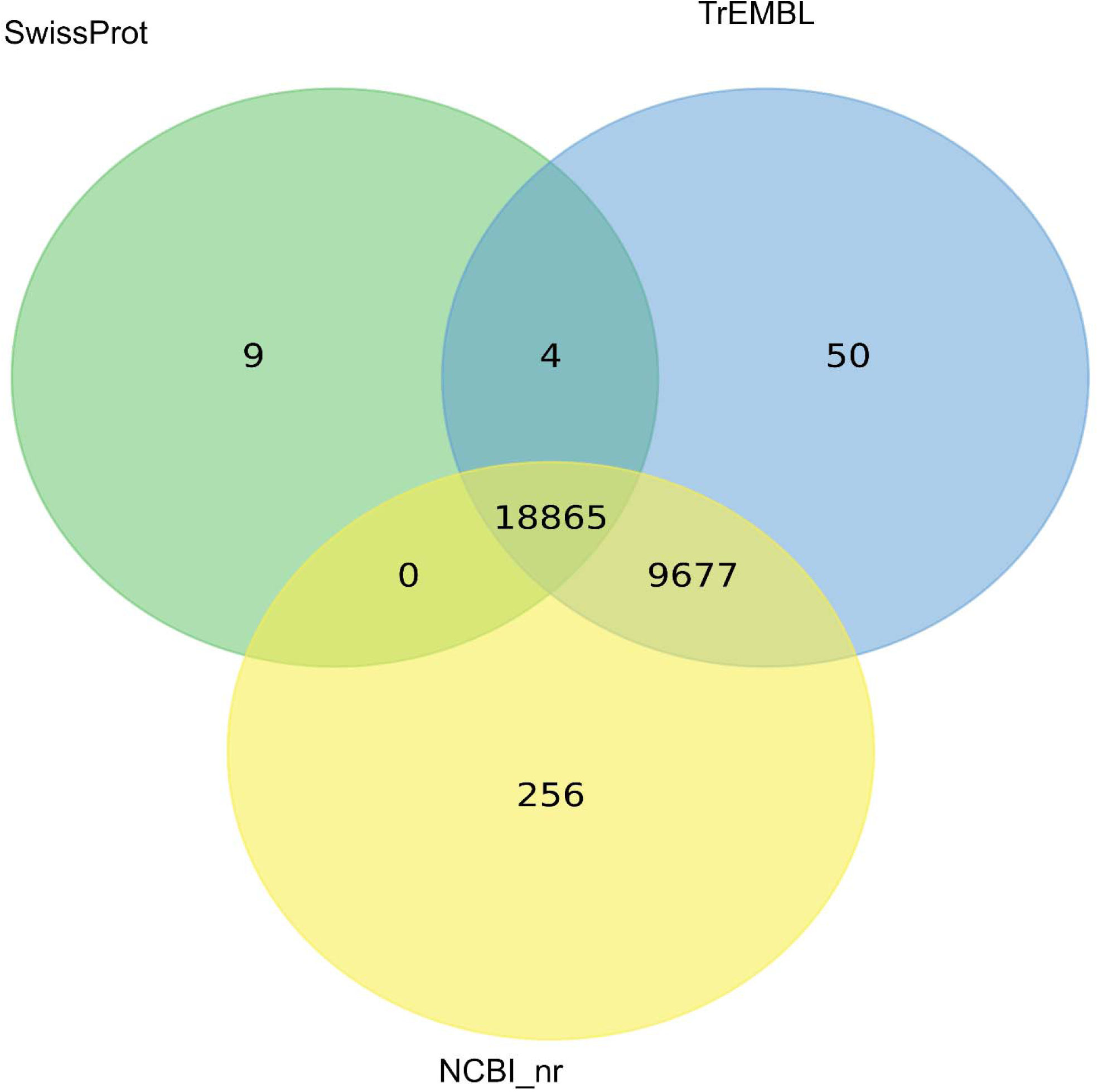
Comparison between the functional annotation of protein coding genes in *A. rufa* using SwissProt, TrEMBL and NCBI_nr. Venn diagram represents total number of unique and common annotated queries genes found in the three databases.

A KEGG-based functional annotation mapped 12377 of our predicted protein-coding genes to their representative functional KEGG orthologs (KO) genes. Data file S1 summarizes the overall KEGG functional annotation. The largest number of genes were mapped to genetic information processing (2968 genes), environmental information processing (1785 genes), and molecular function-related signaling and cellular processes (1664 genes). The top five KEGG metabolic pathways were carbohydrate metabolism (342 genes), lipid metabolism (306 genes), glycan biosynthesis and metabolism (220 genes), amino acid metabolism (180 genes), and nucleotide metabolism (148 genes) (Data file S1, Figure S1).

We reported the annotation profile of non-coding RNAs (ncRNA) in the assembled genome with respect to their Rfam families. We identified 305 transfer RNA (tRNA) through tRNAScan. Additionally, employing Infernal we were able to identify 246 micro-RNA (mRNA), 135 ribosomal RNA (rRNA) and 315 small nuclear RNA (snRNA) genes (Supplementary data file S1).

### Synteny analysis between the genome structures of *A. rufa, C. japonica* and *G. gallus*

*A. rufa, C. japonica, and G. gallus* exhibit a shared karyotype of n = 39 chromosomes^21^. This karyotype similarity motivated us to compare the sequence of the largest 23 scaffolds of *A. rufa* (containing at least 90% of the assembled genome) across the three species. Figure 4 highlights significant synteny regions across the three genomes. Scaffolds 2 and 5 of *A. rufa* align with chromosome 1 in the two other species. Similarly, scaffolds 1, 3 and 4 respectively align to chromosomes 2, 4 and 3 of both birds. Furthermore, *A. rufa* scaffolds 6 and 10 display near complete synteny with *C. japonica*’s sex chromosome Z, while scaffold 10 showing synteny with *G. gallus*’ Z chromosome. Scaffolds 7 and 15 of *A. rufa* display considerable synteny with chromosome 5 of the other birds. The remaining 14 *A. rufa* scaffolds exhibit strong synteny with individual chromosomes of the other two bird species. It is worth noting that twelve micro chromosomes from *C. japonica* and 20 micro chromosomes from *G. gallus* did not exhibit significant homology with any of the assembled *A. rufa* scaffolds.

**Figure 4:**
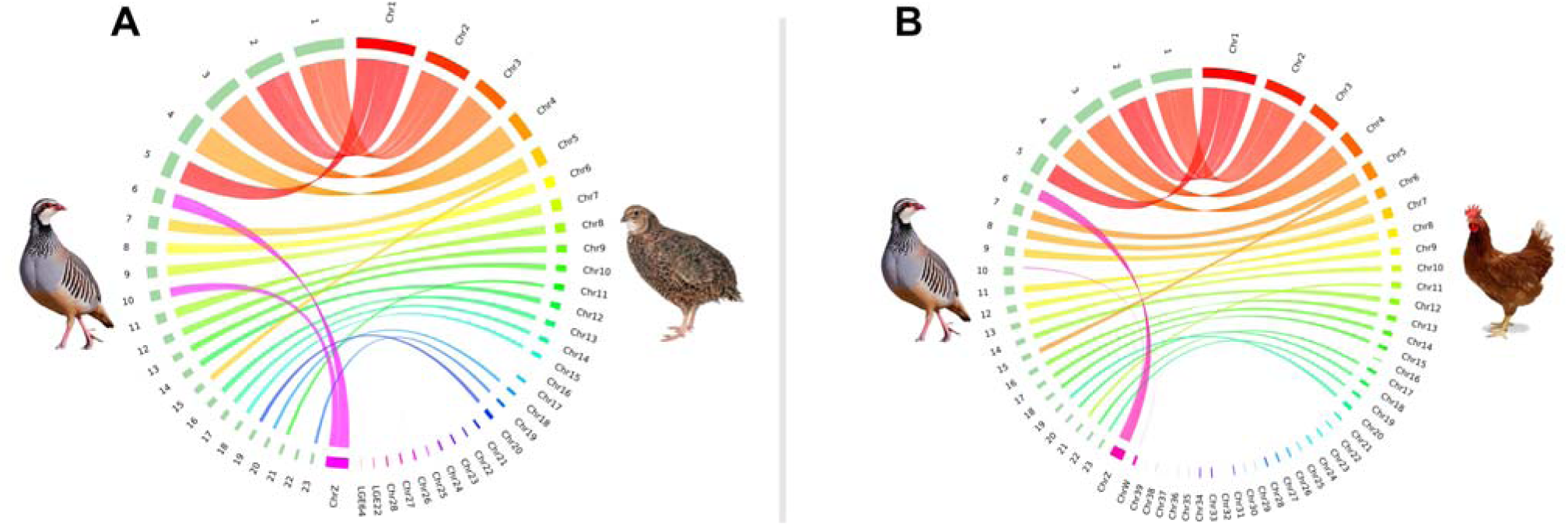
Circus plots comparing sequence homology between the largest 23 *A. rufa* scaffolds and the reference chromosomes of A – *C. japonica*, and B – *G. gallus*. Each line within the circle represents 10kb of sequence homology. Chromosomes are color coded to facilitate visualizing the synteny regions between *A. rufa* and the other two birds.

### Pairwise analysis of the chromosomal rearrangements between *A. rufa and C. japonica* or *G. gallus*

The scaffold-to-chromosome alignments revealed significant large-scale genomic rearrangements between *A. rufa* and both *C. japonica* and *G. gallus* genomes (Figure S2, Table S6). Scaffold 2 exhibits a small 2.52 Mb inversion within the 105.87 to 108.07 Mb region of *C. japonica*’s chromosome 1. Scaffold 5 presents two similar-sized inversions, occurring at regions 19.06 to 20.95 Mb and 50.02 to 57.48 Mb of chromosome 1. Scaffold 1 displays a substantial inversion in its center relative to the centromeric region of *C. japonica*’s chromosome 2 (42.9 to 77.77 Mb). Scaffold 3 features two inversions near one of its ends compared to chromosome 3. Similarly, scaffolds 4 and 18 exhibit inversions when aligned to chromosomes 4 and 15, respectively.

Pairwise alignment of our scaffolds with *G. gallus* chromosomes unveiled repeated inversions, particularly at telomeric regions. Noteworthy instances include scaffold 4, which displayed two inversions totaling 4.37 Mb within regions 1.76 to 4.29 Mb and 0.02 to 1.77 Mb of chromosome 4. Similarly, scaffold 8 exhibited three inversions totaling 3.23 Mb between regions 7.3–8.46 Mb, 9.97– 11.06 Mb, and 11.81–12.72 Mb, aligning with chromosome 6 of the *G. gallus* genome. Additionally, scaffold 11 featured a substantial 8.35 Mb inversion relative to the 0.06 to 8.07 Mb region of chromosome 8.

Overall, these results suggest that *A. rufa*’s genome is more similar to that of *C. japonica* than to that of *G. gallus,* indicating a closer evolutionary relationship between *A. rufa* and *C. japonica* when compared to the *G. gallus*. The similarities in genomic structures and rearrangements between *A. rufa* and *C. japonica* genomes imply a closer evolutionary proximity between the two birds with respect to *G. gallus*.

### Comparative proteome of A. rufa, C. japonica, G gallus, and M. gallopavo

A comparison of the protein coding genes between the four species reveals 12272 shared orthologous gene families (Figure 5A). We have also identified 331 gene families that are exclusive to *A. rufa*, containing 4949 genes. Among these, 556 genes could be functionally annotated using GO at a coarse biological process level (Data file S1, summarized in Figure 5B). Among the gene families linked to more specific GO components, 6 genes were associated with membranes, and 19 genes were associated with structural, binding, or catalytic activities. We remark that the set of genes unique to *A. rufa* (Figure 5C) is notably enriched in genes related to viral processes (24 genes) and signal transduction (21 genes).

**Figure 5:**
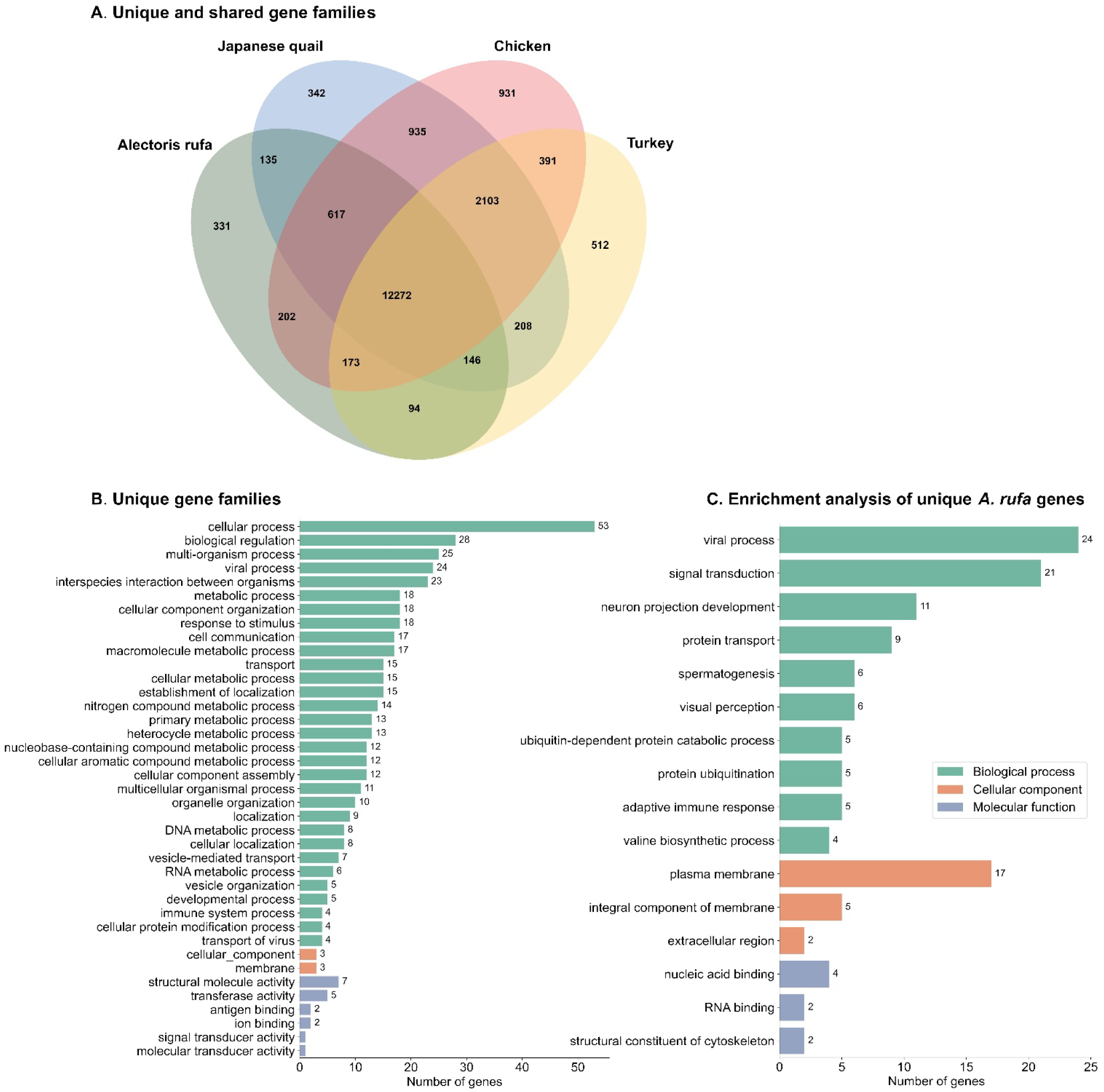
Functional comparison of *A. rufa*’s proteome to that of *C. japonica*, *G. gallus*, and *M. gallopavo*. A – Comparison of orthologous gene families between *A. rufa, C. japonica, G. gallus* and *M. gallopavo*. B – Generic GO enrichment terms for gene families that are unique to *A. rufa.* C – Specific GO enrichment terms for gene families that are unique to *A. rufa*. Only GO categories that are associated to more than one gene are included in panels B and C.

### Phylogenetic analysis of *A. rufa* within the *Galliformes* clade

The phylogenetic tree (Figure 6), constructed through the alignment of single-copy genes across twelve genomes (Table S3), unveils pivotal points in evolutionary history measured in million years ago (Mya). The divergence between birds and reptiles occurred roughly 300 Mya. Anseriformes and Galliformes parted ways around 73 Mya, with the Guinea fowl diverging from the main Galliformes lineage approximately 61 Mya. The clade containing *G. gallus*, *C. japonica* and *A. rufa* separated from the rest of the Galliformes approximately 53 Mya, with their last common ancestor estimated at 37.2 Mya. The divergence between *C. japonica* and *A. rufa* happened approximately 29.9 Mya. Notably, these predicted divergence timelines align with findings from chromosomal synteny analysis (Figure 4).

**Figure 6:**
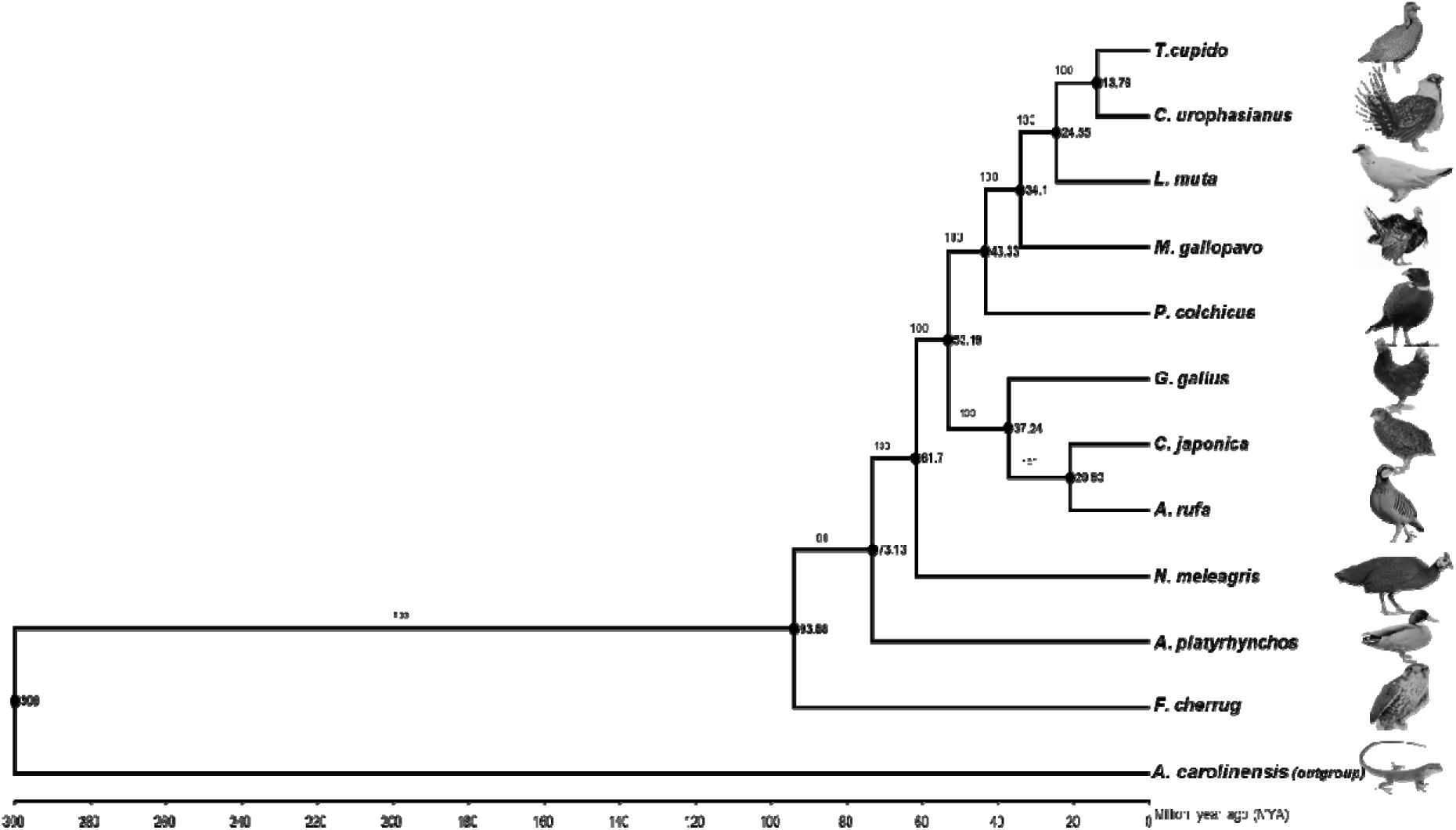
Whole genome based phylogenetic analysis of *A. rufa*. The phylogenetic tree was reconstructed from concatenated single-orthologous genes of 11 birds using IQTREE. Anolis carolinensis (lizard) was used as an outgroup. *A. rufa* is closer to *C. japonica* than to *G. gallus*. Numbers at each node represent the estimated divergence time in million years. Only branching points supported by 100% of bootstrapped trees are shown.

## DISCUSSION AND CONCLUSION

We achieved a highly contiguous genome assembly for *A. rufa* by integrating accurate short reads from Illumina sequencing with lower accuracy ultra-long reads from the Oxford Nanopore Technology (ONT). The resulting assembly is nearly chromosome-level, comprising 115 DNA scaffolds, with a L90 of 23. Our approach demonstrates superior contiguity and scaffolding accuracy compared to previous assemblies relying solely on either short-read ^19^ or long-read data (accession number GCA_947331505.1 at the NCBI), further validating the efficacy of the combined sequencing approach for *de novo* genome assembly in non-model organisms. Additionally, the sequences from sixty *A. rufa* individuals provides a valuable reference for future genetic studies characterizing genome size, ploidy, and heterozygosity rates in different *A. rufa* populations. Our assembly contributes to the collection of avian genomes and highlights the effectiveness of integrating long-read and high-quality short-read data from Illumina^10,16,22^.

Notably, the contiguity statistics for the *A. rufa* genome rank among the top three when compared to all fully sequenced bird genomes analyzed (Figure 2A). Assessment of assembly completeness using BUSCO ^23^ places our assembly in the top four, with the highest number of single copy orthologous genes identified (Figure 2B). We note that the BUSCO assessment of the gene annotation using BUSCOs is lower (87.9% completion rate, Figure 1D) that that for the assembled genome. This discrepancy between the recovered BUSCO genes and the annotated gene set is consistently observed in similar cases ^24^.The highly contiguous assembly facilitated a comprehensive genome annotation by leveraging diverse functionally annotated sequence databases and pre-existing transcriptomic data. As a result, we could use sequence homology to assign biological function for over 95% of all genes identified in our assembly. Overall, our assessment of the annotation quality using RNA sequencing data showed a complete alignment with gene models, with no missed single exons (Table S5). Out of 30236 genes only 8509 were confirmed in terms of gene position and expression by the mapped full transcript. The existing transcriptomic data aligns well with gene model annotation, but numerous predicted genes lack validation due to the limited availability of transcriptomic data for only spleen and skin. Thus, our genome annotation could very like be improved when transcriptomic data for additional tissues becomes available.

Our thorough validation revealed that this difference primarily arises from shortcomings in gene prediction tools and annotation algorithms, rather than the process of masking DNA transposable elements in the genome. To enhance genome annotation accuracy, future improvements should involve integrating transcriptomic data from additional organs or tissue. The genomic annotation of TEs in the *A. rufa* genome shows a high abundance of LINE (7.74% of the genome) and LTR repeat elements (1.76% of the genome). These numbers are higher than those found in the genomes of *G. gallus*^25^ (∼3% LINE and ∼0.5% LTR) and *C. japonica*^26^ (∼5.60% LINE, and ∼0.60% LTR, Table S6). The genomes of *C. californica*^27^ and *C. virginianus* both have a percentage of LINE (6.9% and 5.6% respectively) and LTR (5.6% and 1.73% respectively) more similar to that found in *A. rufa*. Given that transposable elements were found to influence color ^28,29^ in insects, mammals, birds and other vertebrates, a future analysis of the genome should reveal if any genes involved in color determination are found within regions containing TEs.

Notably, 13% of the annotated genes are associated with metabolic functions, while 11% are involved in processing environmental information, including 9% dedicated to signal transduction tasks. The distribution of tRNA genes in the *A. rufa* genome indicates that 11% code for alanine-tRNA and 9% for serine-tRNA (Supplementary data file S1, Supplementary Table S7). Our gene enrichment analysis suggests that *A. rufa* evolved a distinct set of regulatory genes and viral response proteins, likely shaped by species-specific infections and pressures. These findings align with previous transcriptomic analyses that highlighted heightened immune responses in the *A. rufa*^30^.

*A. rufa*, *C. japonica*, and *G. gallus* (*Phasianidae* family) have a diploid genome with 78 chromosomes while *C. virginianus* or *C. californica (Odontophoridae family)* have 82 and 84, respectively. A structural genomic comparison between the three *Phasianidae* birds using chromosome-mapping approaches shows that chromosomal coverage and synteny is stronger between *A. rufa* and *C. japonica* than between *A. rufa* and *G. gallus*. Still, several chromosomal inversions (Figure S2, Table S6) highlight that the divergence between *A. rufa* and *C. japonica* is far from recent. For example, aligning scaffold 1 of *A. rufa* to chromosome 2 of *C. japonica* reveals an inversion that contains the centromeric region of the chromosome. Our sequence comparison between scaffold 4 of *A. rufa* and chromosome 4 of *G. gallus* reveals another centromeric inversion. This inversion had been previously reported by ^4^ based on cytogenetic analysis. Still, we note that inversions detected close to centromeres and telomeres may result from mis-assemblies, due to the higher DNA repeat content in those genomic regions. However, because the borders for the inversions we find in our assembly are well outside of the repetitive regions themselves, we strongly believe that they are not an artefact. In fact, they are also consistent with similar massive inversions observed within independent populations of *C. coturnix*, another quail species, and associated to an expansion of phenotypic diversity between populations ^31^. These and other observations in our analysis emphasize the potential interest of future research focusing on *A. rufa*’s evolutionary chromosomal rearrangements. However, a lack of a genetic map or higher-resolution optical map data for this species currently limits our ability to make definitive conclusions about those structural rearrangements at scaffold assembly-level.

The genome assembly provided here is also of interest for phylogenetic studies. Phylogeny proposes an evolutionary tree that aids our comprehension of species divergence over time, drawing upon evidence from paleontology, biogeography, and genetics ^32–34^. The integration of both mitochondrial and nuclear markers significantly advanced the accuracy of those studies ^35^. Combining those markers with estimates of divergence time gleaned from fossil records and genetic clocks has produced robust phylogenetic trees that can be used to generate strong hypotheses about speciation events ^36–38^. Notably, within the bird group and the order Galliformes, a variety of studies proposed multiple and often coinciding clade formation hypotheses ^39,40^. Leveraging assemblies to create genome wide alignments from which to create phylogenetic trees enables a more accurate understanding of the evolutionary history of life.

Overall, our assembly and annotation provide a significant contribution towards a reference genome of the red-legged partridge, which will aid in developing genetics applied to phylogeny, zoology, demography, and ecology of the species. This near-chromosome assembly provides a foundation upon which to anchor future comparative genomics research between different *A. rufa* populations and across *Phasianidae* species. It is a valuable resource, potentially enabling the development of more effective strategies for management and conservation of *A. rufa* and wildlife.

## METHODS

### Genome sequencing data

Total DNA was obtained from the muscle of sixty frozen *A. rufa* individuals (30 wild birds and 30 farm birds) for whole-genome sequencing on the NovaSeq6000 Illumina platform producing short paired-end reads with a read length of 151 bp as described in ^2^. Additionally, we extracted high molecular weight (HMW) DNA from the blood of two live individuals for library preparation with the genomic DNA sequencing kit of Oxford Nanopore technology (ONT) and then sequenced the libraries using a GridION platform.

### Processing sequencing data

The Illumina sequencing yielded an average of 218 million raw reads per individual, with an average depth sequencing of 32X per sample. We assessed the quality of those reads using FastQC^41^. The per-base quality scores were consistently high across all samples, and no adaptor content within the reads was found. Thus, it was determined that additional cleaning and adapter removal procedures were unnecessary.

We generated 2 million raw ultra-long reads of the Oxford Nanopore Technology (ONT), yielding 48 Gb with an average read length of 20.68 Kb (Table S8). We used Porechop V.0.2.4 ^42^ with default parameters in order to scan for known Nanopore adapters and to trim them out of the long reads, ensuring a high-quality dataset, free of adaptor contamination. We assessed the quality of this dataset using NanoPlot v1.40.2 (part of the NanoPack software suite) ^43^. We then used Filtlong ^44^ to split the reads into two subsets applying different criteria. For the first subset, we prioritized read length over average read quality, selecting a coverage depth of 40x (--min_length 15kb -t 40 Gb). For the second subset, we prioritized average read quality over read length, generating a coverage depth of 20x (-- min_mean_q 12 -t 20 Gb). By using these two different subsets, we aimed at improving genome contiguity while also correcting structural errors, ensuring a more reliable and accurate analysis of the sequencing data.

### Genome size estimation

We used a 21-mer-based approach in Jellyfish v.2.2.10 ^45^ to estimate *k*-mer histogram frequencies from the Illumina paired-end sequencing data of each of the sixty individual birds. The output of Jellyfish was then used in GenomeScope2 ^46^ to estimate genome size and heterozygosity level for the genome of each bird. In addition to the genome profiling with genomescope2 on those short reads, Smudgeplot^46^ was used to estimate the ploidy level using Nanopore long reads sequencing data.

### Hybrid genome assembly

Figure S3 summarizes the pipeline we employed to create a *de novo* assembly for the genome of *A. rufa*, using a hybrid approach. The raw ONT long-reads were assembled *de novo* with Flye ^47^, Canu ^48^, Wtdbg2 ^49^, and NextDenovo v2.2.4 ^50^. In order to select the best primary assembly for further procedures we compared the performance of the four assemblers. We used QUAST v5.2.0 ^51^ to calculate the contiguity statistics of each assembly statistics and the aves_odb10 dataset of Benchmarking Universal Single-Copy Orthologs (BUSCO) v.5.4.3 ^23^ to assessed their completeness. Based on these numbers we chose the NextDenovo contig-level assembly for further improvement.

We combined long and short read information to improve the contig-level assembly. This hybrid approach comprised two main steps to enhance the assembly quality. First, we mapped the subset of long-read ONT with 40x and min length size of 15 kb, to the contig-level assembly using minimap2 ^52^. This alignment was then input into RACON v.1.5.1 ^53^ for one polishing iteration, improving the contiguity of the contig-level assembly by correcting several structural assembly errors. Then, we aligned the short reads from the sixty *A. rufa* individuals to the RACON-improved assembly using BWA-mem2 v2.2.1^54^. This alignment was the input for Polypolish v0.5.0 ^55^, which we used to polish the RACON-improved draft and fix small SNPs and indels, leveraging the high coverage of short-reads to generate a high-quality consensus assembly that represents the genetic diversity of *A. rufa*’s genome. We completed the scaffolding of the assembly using the REDUNDANS pipeline ^56^. We ran this pipeline using the "--non-reduction --nogapclosing" parameters to enhance genome scaffolding, using a subset of both long and short reads in combination from the original raw reads. We combined the subset of accurate long-read ONT with 20X sequencing depth and the short reads of the two animals with the highest genome coverage, aiming at improving scaffold accuracy. The final scaffold-level assembly served as the foundation for downstream genome annotation and comparative analysis.

### Genome screening for contamination sequences

Before annotating the assembled genome, we conducted a thorough screening process to identify and eliminate any sequences that might be contaminants related to the assembled genome of *A. rufa*. To do this, we employed NCBI’s Foreign Contamination Screening (FCS) tools ^57^ FCS-adapter and FCS-GX. We used FCS-adapter to detect adaptors and vectors. We used FCS-GX to identify foreign DNA contamination sequences by aligning our assembly against the NCBI database of genomes. We ran each of these tools independently using default settings, except for the taxonomic identifier, which was set to be that of *A. rufa* (NCBI: txid 9079). This rigorous screening process helped ensure the integrity of our assembled genome data before proceeding with annotation.

## Genome Annotation

### Annotation of transposable elements

We used EDTA v2.1.1 ^58^ to annotate the DNA transposable elements (TEs) in our assembled genome. EDTA integrates a set of open-source programs for TE annotation based on homology and/or *ab initio* search methods. We used two independent data sets to increase the accuracy of EDTA annotation. First, we downloaded a curated library from the gold-standard database of repetitive sequences msRepDB ^59^. This library contained DNA transposable sequences for six closely related bird species (*Alectoris Barbara*, *Alectoris philbyi*, *Alectoris melanocephala*, *Coturnix japonica*, *Meleagris gallopavo*, and *Gallus gallus*; Table S9). Then, the CDS sequences of *G. gallus* were downloaded from ENSEMBL release 109 ^60^, to remove gene-related sequences. In parallel, we used RepeatModeler V2.0.3 ^61^ with default parameters for additional *ab initio* annotation of repetitive elements. Finally, we combined the results from EDTA and RepeatModeler to build a non-redundant library of repetitive elements using our in-house scripts. This custom TEs library was used as input to the RepeatMasker v4.1.4 ^20^ ^62^ for soft masking of the *A. rufa* genome. We ran RepeatMasker using the following parameters: “-e ncbi -gff -s -a -inv -no_is -norna -xsmall -nolow -div 40”, against the Dfam ^63^ and RepBase update 18. We then used the soft-masked genome for further annotation.

### Divergence distribution of transposable element

We analyzed RepeatMasker’s alignment output file using the parseRM.pl script v5.8.2 available at ^62^. We determined the percentage of divergence from the consensus for each TE fragment, considering the elevated mutation rate at CpG sites and employing the Kimura 2-Parameter divergence metric. This divergence percentage serves as a measure of the age of the TE fragments, as older TE invasions accumulate more mutations. We further categorized TEs fragments by age, organizing them into bins of 1 million years, based on the substitution rate calculated by parseRM.pl. We then plotted the distribution landscape of TE using a custom R script.

### Gene structure annotation

We combined three strategies to annotate the protein coding genes in the soft-masked genome: homology-based, transcriptome-based, and *ab initio* predictions:

1- We ran Miniprot v.0.10-r225 ^64^ for homology-based gene prediction. A dataset comprising 3,044546 protein sequences was generated. These sequences were obtained from the NCBI reference sequence of proteins (accessed on April 15, 2023). Specifically, we focused on the Aves NCBI:txid8782 lineage to ensure retrieval of only avian proteins. Additional details about this dataset can be found in Table S10.
2- We ran PASApipeline v.2.5.3 ^65^ to perform gene prediction based on the transcriptional evidence provided by the transcriptome assembly of *A. rufa* published in 2017 ^30^.
3- For *ab initio* gene prediction, we ran BRAKER2 v.2.1.6 ^66^, training it with the same dataset we used for Miniprot.

The annotation results of the three approaches were then combined using EVidenceModeler v.2.1.0 ^65^ to produce a more reliable and consensus gene set model of the assembled *A.rufa* genome.

### Non-coding RNA gene annotation

We also annotated non-coding RNA genes (ncRNAs) in our genome assembly. We used tRNAscan-SE2 v.2.0.11 ^67^ to identify transfer RNAs (tRNAs). Infernal v.1.1.4 ^68^ was run to identify microRNAs (miRNAs), ribosomal RNAs (rRNAs) and small nuclear RNAs (snRNAs), based on the Rfam database (release 14.0) ^69^.

### Functional annotation

We assigned functions to the predicted gene models combining various approaches. A standard e-value cutoff of 1e-6 was applied for sequence comparisons, unless otherwise specified. Initially, we utilized eggnog-mapper v.2.1.10 ^70^ against the eggNOG database ^71^ to assign Gene Ontology terms. Subsequently, Blastp v2.12.0+ was employed against SwissProt, TrEMBL ^72^, and NCBI NR ^73^ databases for homology-based functional annotation (All the public protein databases mentioned above were accessed on April 15, 2023). Priority was given to matches with over 95% identity from SwissProt and TrEMBL, as annotation of proteins in these databases is more reliable because of manual curation. The resulting functional annotations were combined with InterProScan v5.64.-96.0 ^74^. InterProScan identified protein domains, families, and superfamilies in annotated protein-coding genes using specified parameters “-m diamond --sensmode fast --go_evidence non-electronic”. KofamKOALA v1.3.0 ^75^ assigned KEGG orthologs (KO) and pathways with an e-value cutoff of 1e-9. Functional annotations from the Uniprot database (minimum Blastp homologue identity match of 95%) were integrated into the final genome annotation file using the GAG tool v2.0.1 ^76^.

### Quality assessment of genome assembly and annotation

We used QUAST v5.2.0 to calculate correctness and contiguity metrics for the genome assembly. We used BUSCO against the aves_odb10 v2019-11-20 database to assess both the completeness of the assembly and of the annotation of structurally predicted protein-coding genes.

### Quality assessment of the genome annotation using RNA sequencing

As part of evaluating the accuracy of gene model annotations, we downloaded a set of RNA sequencing transcriptome from the spleen and the skin of the red-legged partridge (*A.rufa*) that are deposited in the NCBI SRA database (Table S11). We employed the STAR aligner tool v2.7.10b ^77^ for mapping these reads to the soft-masked assembly version. Subsequently, each sample underwent transcript-assembly guided using Stringtie v2.2.1 reference-guided assembler of transcripts. The spliced transcripts from all samples were combined using Stringtie into a one master list of transcripts, the output of Stringtie was retrieved in the GFF file format for suitable downstream analysis. Next, we used the GffCompare v2.12.6 tool ^78^ to compare this list of annotated transcripts with respect to the final annotated gene set model. This comparison helped determine the number of new spliced transcripts that were not previously identified in our gene set, contributing to our assessment of gene annotation quality.

### Comparison to the reference genomes of *C. japonica* and *G. gallus*

We used MUMMER v.4 ^79^ to perform whole-genome alignment between our assembly and the fully sequenced genomes of *C. japonica* and *G. gallus*. The genome pairwise alignment results and synteny blocks of 10kb were visualized with DOT-PLOT viewer ^80^ and Circos v.0.69-8 ^81^.

### Gene family analysis

We used the OrthoVenn3 pipeline ^82^ to compare gene families between *A. rufa*, *C. japonica*, *G. gallus*, and *G. pavo*. In brief, Orthofinder ^83^ was used to compute the orthologs between the species of interest and to cluster gene families based on GO functional annotation categories. Additionally, we also used the pipeline to automatically conduct GO terms enrichment analysis by considering the evolutionary relationship between the four species.

### Phylogenomic analysis and divergence time tree building

We performed phylogenetic analysis to infer the divergence time of *A. rufa* with respect to other birds with fully sequenced genomes within the *Galliformes* order, in a way that is similar to previous reports ^14,84,85^. In addition to *A. rufa*, we included 8 genome protein sequences of Galliformes species, of which 7 species belong to Phasianidae family and one to Numididae family (*Numida meleagris*). We also included one genome from the *Anseriformes* order (*Anas platyrhynchos*), and another bird species for the Falconiformes order (*Falco cherrug*). As an outgroup we used *Anolis carolinensis* ^86^ from the Reptilia class.

Genome assemblies for the birds and outgroup (*Anolis carolinensis*) from Reptilia were downloaded from the NCBI. Detailed information about those species can be found in Table S3. We started by using the aves_odb10 database of the BUSCO tool ^23^ to identify shared single-copy genes in the twelve analyzed genomes. The aves_odb10 database contains 8338 genes. We used the custom Python script available at https://github.com/jamiemcg/BUSCO_phylogenomics.git to extract the shared single-copy orthologs common to all species. We independently created multiple alignments for each of the orthologs common to all species, using MUSCLE ^87^. We concatenated the resulting multiple alignments to create a supermatrix alignment. To ensure alignment quality, we applied trimAI ^88^ and removed unreliable aligned sites and gaps.

Subsequently, a phylogenetic tree was constructed using IQTREE v ^89^, incorporating 1000 bootstrap replicates. The best model for tree construction was determined using the ModelFinder package ^85^ from the IQTREE suite. To estimate the divergence time of *A. rufa* in relation to the other birds, we used the MCMCtree tool from the PAML package ^90^. MCMCtree used the phylogenetic tree generated by IQTREE and the alignment file to achieve reliable divergence time estimation, minimizing potential outliers. Three fossil calibration times from the TimeTree5 ^91^ were employed for divergence estimation: *G. gallus* – *C. japonica* (≈ 32.9 - 46.1 Million years ago (Mya)), Numida-Mallards (≈ 72.5 - 85.4 Mya), and the divergence time between birds and reptiles (≈300 – 250 Mya) ^92^. We ran MCMCTREE on protein-coding sequences, sampling 20000 times with a sampling frequency of 10, following a burn-in of 2000 iterations. We used default parameter for the other settings.

Table S12 summarizes all bioinformatics pipelines, tools versions, and settings used during the genome assembly and annotation process and other related analysis used in this work.

## Data Availability

The Nanopore raw read data are available via ENA (Bioproject accession PRJEB67643, Biosample: ERS16499794, ERS16499793, ERS16499792, ERS16499791, ERS16499790). Genome assembly sequence and annotation are available via NCBI under the Bioproject accession PRJNA1050768 and ENA accession number ERZ22146366. This work was performed under the scope of the Catalan Biogenome Project (CBP).

## Code Availability

The source code and relevant data files used to generate each figure in this manuscript are available on the GitHub repository page of the Systems Biology and Statistical Methods Group at https://github.com/BioModelLab/A.rufa_genome.git

## Acknowledgements/Funding

Fundação para a Ciência e a Tecnologia (FCT), I.P., is acknowledged for funding A. Usié through Contrato–Programa (CEECINST/00100/2021/CP2774/CT0001) and for Projects UIDB/05183/2020 to Mediterranean Institute for Agriculture, Environment and Development (MED), and LA/P/0121/2020 to CHANGE—Global Change and Sustainability Institute. We are grateful for the contributions made by the Melgarejo family, Patricia, Luis and Ivan Maldonado and Tom Gullick. Thanks also to the “Las Ensanchas” staff, especially the game keepers, the Barranquero family and collaborators, the members of the Tom Gullick hunting team in Campo de Montiel and around the world, Federación de Caza de Castilla y León, Delegación Burgalesa, MUTUASPORT, and Real Federación Española de Caza (RFEC). Carolina Ponz helped in sampling. The anonymous referees provided valuable comments that improved the manuscript. Fundación Universitat Rovira i Virgili funded the sequencing (grant no. 2060-398-454-455).

