## Supplementary for "An almost chromosome-level assembly and annotation of the *Alectoris rufa* genome"

### Supplementary Materials

#### 1. Supplementary tables

Table S1: Statistics of assembler tools used to draft the primary contig-assembly level of the *A. rufa* genome and their BUSCO completeness.

| Contiguity / completeness metrics | Assembler tools | | | |
| --- | --- | --- | --- | --- |
|  | **Canu** | **Wtdbg2** | **Flye** | **nextDenovo** |
| Number of contigs/scaffolds | 2356 | 1313 | 537 | 116 |
| Assembly size (Gb) | 1.28 | 1.04 | 1.03 | 1.03 |
| N50 (Mb) | 3 | 27 | 34 | 74 |
| N90 (Mb) | 0.14 | 4 | 5 | 10 |
| Average length of contigs (Mb) | 0.5 | 0.7 | 1.9 | 8 |
| Largest contig (Mb) | 17 | 132 | 113 | 149 |
| GC % | 41.94 | 42.30 | 42.22 | 42.15 |
| Run time (hours) | 535 | 16 | 20 | 8 |
| Complete BUSCO | 7411 | 7118 | 7611 | 7586 |
| Complete and single-copy BUSCO | 6610 | 7095 | 7579 | 7555 |
| Complete and duplicated BUSCO | 801 | 23 | 32 | 31 |
| Fragmented BUSCO | 288 | 334 | 487 | 525 |
| Missing BUSCO | 639 | 886 | 478 | 525 |
| Total BUSCO groups searched | 8338 | | | |

Table S2: Completeness assessment using BUSCO and Mercury-kmer* consensus error.

|  | Primary contigs | Polished contigs 1 (Long-reads) | Polished contigs 2 (Short-reads) | Scaffolds |
| --- | --- | --- | --- | --- |
| BUSCO completeness | 91% (7586) | 90.8% (7579) | 96.9% (8078) | 96.9% (8078) |
| Complete and single-copy BUSCO | 90.6% (7555) | 90.5% (7550) | 96.6% (8053) | 96.6% (8051) |
| Complete and duplicated BUSCO | 0.4% (31) | 0.3% (29) | 0.3% (25) | 0.3% (21) |
| Fragmented BUSCO | 2.7% (227) | 3.1% (258) | 0.5% (39) | 0.5% (39) |
| Missing BUSCO | 6.3% (525) | 6.1% (501) | 2.6% (221) | 2.6% (227) |
| Total searched BUSCO | 8338 | | | |
| Mercury k-mer completeness | | | | |
| QC | 41.1 | 38.7 | 36.3 | 36.3 |
| ERROR | 7.7e-05 | 1.3 e-04 | 2.4 e-04 | 2.4 e-04 |

*The k-mer completeness is calculated as the fraction of reliable k-mers in the read set that are also having to be found in the assembly.

Table S3: Genome assemblies’ species of Aves and one reptile used to construct the phylogeny tree and gene family in relation the red-legged partridge.

| NCBI genome assemblies | Species | Common name | Reference |
| --- | --- | --- | --- |
| GCF_001577835.2 | *Coturnix japonica* | Japanese quail | (2) |
| GCF_016699485.2 | *Gallus gallus* | Chicken | (3) |
| GCF_000146605.3 | *Meleagris gallopavo* | Turkey | (4) |
| GCF_002078875.1 | *Numida meleagris* | Guineafowl | (5) |
| GCF_004143745.1 | *Phasianus colchicus* | Common pheasant | (6) |
| GCF_000337975.1 | *Falco cherrug* | Saker falco | (7) |
| GCF_023343835.1 | *Lagopus muta* | Rock ptarmigan | (8) |
| GCF_019232065.1 | *Centrocercus urophasianus* | Greater sage-grouse | (9) |
| GCF_026119805.1 | *Tympanuchus pallidicinctus* | Lesser prairie chicken | (10) |
| GCF_015476345.1 | *Anas platyrhynchos* | Mallard | (11) |
| GCF_000090745.1 | *Anolis carolinensis* (Reptilia) | Green anoles | (12) |
| GCA_008692595.2 | *Colinus virginianus* | Virginia quail | (13) |

Table S4: The NCBI FCS_GX screening contamination summary, assessing the genome diversity, and potential contamination.

| 1. FCS_GX summary: | | |
| --- | --- | --- |
| Putative contaminant divs: | 0 | |
| Asserted div: | birds | |
| Primary divs: | birds,reptiles,vertebrates | |
| Aggregate coverage: | 96% | |
| Minimum pct-coverage: | 20 | |
| Putative contaminant divs: | 0 | |
| 2. FCS_GX contamination summary: | | |
|  | sequences | Bases |
| Total | 0 | 0 |
| 3. FCS_GX action summary: | | |
|  | Sequences | Bases |
| Total | 0 | 0 |
| Excluded | 0 | 0 |

Table S5: Overall supported gene model annotation from RNAs mapped transcript.

| Reference gene model | 30236 in 30236 loci* |
| --- | --- |
| Transcripts | 60826 in 31154 loci (~2.0 transcripts per locus) |
| Matching transcripts | 8509 |
| Matching loci | 8509 |
| Missed exons | 0/30236 (0.0%) |
| Novel exons: | 9510/256604 (3.7%) |
| Novel introns: | 208738/208738 (100.0%) |
| Missed loci: | 0/30236 (0.0%) |
| Novel loci: | 1230/31154 (3.9%) |

Table S6: Large chromosomal rearrangements between *A. rufa,* *C. japonica* and *G. gallus*.

| **Scaffold** | **Start (Mb)** | | **End (Mb)** | **Chr** | **Start (Mb)** | **End (Mb)** | **Description** |
| --- | --- | --- | --- | --- | --- | --- | --- |
| A. ***rufa* vs *C. japonica*** | | | | | | | |
| 1 | 49.26 | 89.17 | | 2 | 42.90 | 77.77 | Centromere regions (inversion) |
| 2 | 42.62 | 45.14 | | 1 | 105.62 | 108.08 | Small inversion |
| 5 | 51.51 | 63.3 | | 1 | 50.02 | 57.48 | Inversion |
| 5 | 20.4 | 22.5 | | 1 | 19.06 | 20.95 | Small inversion |
| 16 | 6.93 | 14.31 | | 13 | 6.54 | 13.19 | Inversion near to the centromeric regions |
| 18 | 9.20 | 12.99 | | 15 | 7.88 | 11.39 | Inversion near to the telomeric region |
| A. ***rufa* vs *G. gallus*** | | | | | | | |
| 3 | 5.88 | 12.03 | | 3 | 5.89 | 11.87 | Inversion telomeric region |
| 4 | 2.11 | 4.68 | | 4 | 1.76 | 4.29 | Inversion 1 near to the telomeric regions |
| 4 | 0.05 | 1.85 | | 4 | 0.02 | 1.77 | Inversion 2 near to telomeric regions |
| 8 | 9.44 | 10.66 | | 6 | 7.3 | 8.46 | Inversion 1 (telomeric region) |
| 8 | 8.09 | 9.21 | | 6 | 9.97 | 11.06 | Inversion 2 (telomeric region) |
| 8 | 11.29 | 12.18 | | 6 | 11.81 | 12.72 | Inversion 3 (telomeric region) |
| 11 | 2.60 | 10.95 | | 8 | 0.06 | 8.07 | Inversion neat to the telomeric region |

Table S6: TE annotation of the December 2023 NCBI release of *C. japonica*’s genome.

|  | Coturnix japonica 2.1 [GCF_001577835.2] | |
| --- | --- | --- |
|  | Length (bp) occupied | % Of genome |
| SINEs | 1262 | 0.01 |
| LINEs | 51989662 | 5.60 |
| LTR elements | 5446838 | 0.60 |
| DNA transposons | 6323524 | 0.68 |
| Unclassified | 14807631 | 1.40 |
| Total | 78771111 | 8.49 |

Table S7: Annotated tRNAs in the genome of *A. rufa*

| **tRNAs isotype** | **Count** | **Copies per anti-codon** | **Total Percentage** |
| --- | --- | --- | --- |
| Ala | 33 | AGC: 22; CGC: 4; TGC: 7 | 11% |
| Gly | 22 | GCC: 8; CCC: 5, TCC: 9 | 7% |
| Pro | 16 | AGG: 9; GGG: 1; CGG: 3; TGG: 3 | 5% |
| Thr | 13 | AGT: 7; CGT: 2; TGT: 4 | 4% |
| Val | 18 | AAC: 6; GAC: 1; CAC: 8; TAC: 3 | 6% |
| Ser | 28 | AGA: 14; CGA: 3; TGA: 3; GCT: 8 | 9% |
| Arg | 19 | ACG: 6; CCG: 3; TCG: 3; CCT: 3; TCT: 4 | 6% |
| Leu | 22 | AAG: 4; GAG: 1; CAG: 8; TAG: 2; CAA: 5 TAA: 2 | 7% |
| Phe | 10 | AAA: 1; GAA: 9 | 3% |
| Asn | 12 | GTT: 12 | 4% |
| Lys | 11 | CTT: 6; TTT: 5 | 4% |
| Asp | 9 | GTC: 9 | 3% |
| Glu | 15 | CTC: 8; TTC: 7 | 5% |
| His | 8 | ATG: 1; GTG: 7 | 3% |
| Gln | 12 | CTG: 9; TTG: 3 | 4% |
| Ile | 9 | AAT: 6; TAT: 3 | 3% |
| Met/iMet | 17 | CAT: 17 | 6% |
| Tyr | 17 | GTA: 17 | 6% |
| Cys | 12 | GCA: 12 | 4% |
| Trp | 6 | CCA: 6 | 2% |
| SelCys | 2 | TCA: 2 | 1% |
| **Overall annotated tRNA** | | **301** |  |

Table S8: Statistics for the ultra-long ONT reads used to assemble the *A. rufa* genome.

|  | **Original raw read** | **Cleaned reads** |
| --- | --- | --- |
| Number of reads (Million) | 2323535 | 2323182 |
| Average read length (bp) | 20690 | 20688 |
| Read length N50 | 40857 | 40857 |
| Total bases (bp) | 48074728669 | 48063123316 |
| Average read quality | 11.3 | 11.4 |
| Estimated coverage (X) per species | 40 | 40 |

Table S9: Species included for homology-based TEs annotation in *A. rufa*‘s genome.

|  | Taxon ID | genome size (Gb) | Assembly-level | TEs database |
| --- | --- | --- | --- | --- |
| *Gallus gallus* | 9031 | 1.05 | Chromosome | msRepDB |
| *Meleagris gallopavo* | 9103 | 1.1 | Chromosome | msRepDB |
| *Alectoris barbara* | 40177 | NA | NA | msRepDB |
| *Alectoris melanocephala* | 40180 | NA | NA | msRepDB |
| *Alectoris philbyi* | 40181 | NA | NA | msRepDB |
| *Coturnix japonica* | 93934 | 0.927 | Chromosome | msRepDB |
| NA: not available. | | | | |

Table S10: NCBI’s bird protein database statistics.

|  | Aves proteins sequences |
| --- | --- |
| Number of proteins | 3044546 |
| Avg. protein length (aa) | 680 |
| Max. protein length (aa) | 36619 |
| Min. protein length (aa) | 20 |
| aa: stand for amino acid |  |

Table S11: Transcriptome raw reads collected from the NCBI used in the annotation of the *A. rufa* genome.

| BioProject | Accession | Tissues | Read length (bp) | Total-reads | Ended | Reference |
| --- | --- | --- | --- | --- | --- | --- |
| PRJNA268542 | SRR1664666 | skin | 76 | 12366816 | SE | (13) |
|  | SRR1664667 | skin | 76 | 13186804 | SE |  |
|  | SRR1664672 | skin | 75 | 7986047 | SE |  |
|  | SRR1664674 | skin | 75 | 10994222 | SE |  |
|  | SRR1664675 | spleen | 76 | 13398642 | SE |  |
|  | SRR1664677 | spleen | 75 | 12187065 | SE |  |
|  | SRR1664678 | spleen | 76 | 13025606 | SE |  |
|  | SRR1664679 | spleen | 75 | 13705093 | SE |  |

SE: single-end

Table S12: Software versions used in this manuscript.

| **Tool** | **Version** | **Parameters** | **GitHub/ URLs** |
| --- | --- | --- | --- |
| Porechop | 0.2.4 | Default | https://github.com/rrwick/Porechop |
| Filtlong |  | (--min_length 15kb -t 40 Gb)  20x (--min_mean_q 12 -t 20 Gb). | https://github.com/rrwick/Filtlong |
| NanoPlot | v1.40.2 | Default | https://github.com/wdecoster/NanoPlot |
| genomeScope2 | Online | Kmer :21 | http://evidencemodeler.github.io/ |
| Flye | 2.9.1-b1780 | Default | https://github.com/fenderglass/Flye |
| Canu | 2.2 | -ONT | https://github.com/marbl/canu |
| Wtdbg2 |  | Default | https://github.com/ruanjue/wtdbg2 |
| nextDenovo | 2.5.0 | Default | https://github.com/Nextomics/NextDenovo |
| Racon | 1.5.1 | Default | https://github.com/isovic/racon |
| bwa-mem2 | 2.2.1 | Default | https://github.com/bwa-mem2/bwa-mem2 |
| BUSCO | 5.4.3 | -l aves_odb10 | https://busco.ezlab.org/ |
| QUAST | 5.2.0 | --fragmented | http://quast.bioinf.spbau.ru/ |
| EDTA | 2.1.1 | --cds | https://github.com/oushujun/EDTA |
| RepeatModeler | 2.0.3 | Default | http://www.repeatmasker.org/RepeatModeler/ |
| RepeatMasker | 4.1.4 | -lib curatedTEs | http://repeatmasker.org/ |
| BRAKER2 |  | --protein | https://github.com/Gaius-Augustus/BRAKER |
| Minimap2 | 0.10-r225 | -ONT | https://github.com/lh3/minimap2 |
| PASA | 2.5.3 | Default | https://github.com/PASApipeline/PASApipeline/ |
| EVidenceModeler |  | Default | http://evidencemodeler.github.io/ |
| tRNAscan-SE2 | v.2.0.11 | Default | http://lowelab.ucsc.edu/tRNAscan-SE/ |
| NCBI_NR | - | - | ftp://ftp.ncbi.nlm.nih.gov/blast/db/FASTA/nr.gz |
| Blastp | 2.12.0+ | e-value of 1e-6 |  |
| miniprot |  |  | <https://github.com/lh3/miniprot> |
| TimeTree | NA | - | http://repeatmasker.org/ |
| Infernal | 1.1.4 | - | http://eddylab.org/infernal/ |
| GAG tool | 2.0.1 | - | https://github.com/genomeannotation/GAG |

#### 2. Supplementary figures

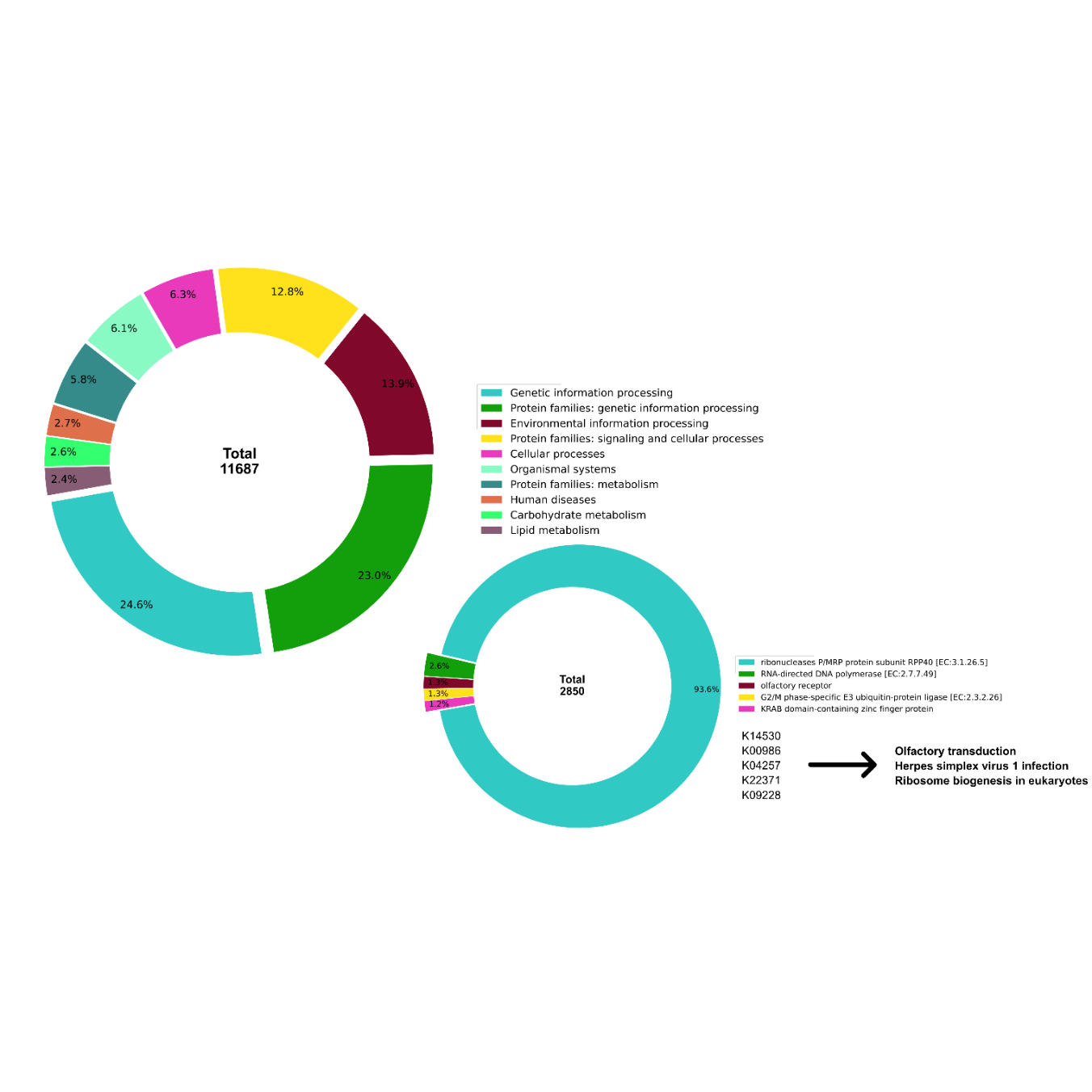

Figure S1: Top 10 KEGG functional categories and Top 5 KEGG cluster of KEGG (KO) orthologs overrepresented in *A. rufa*’s genome.

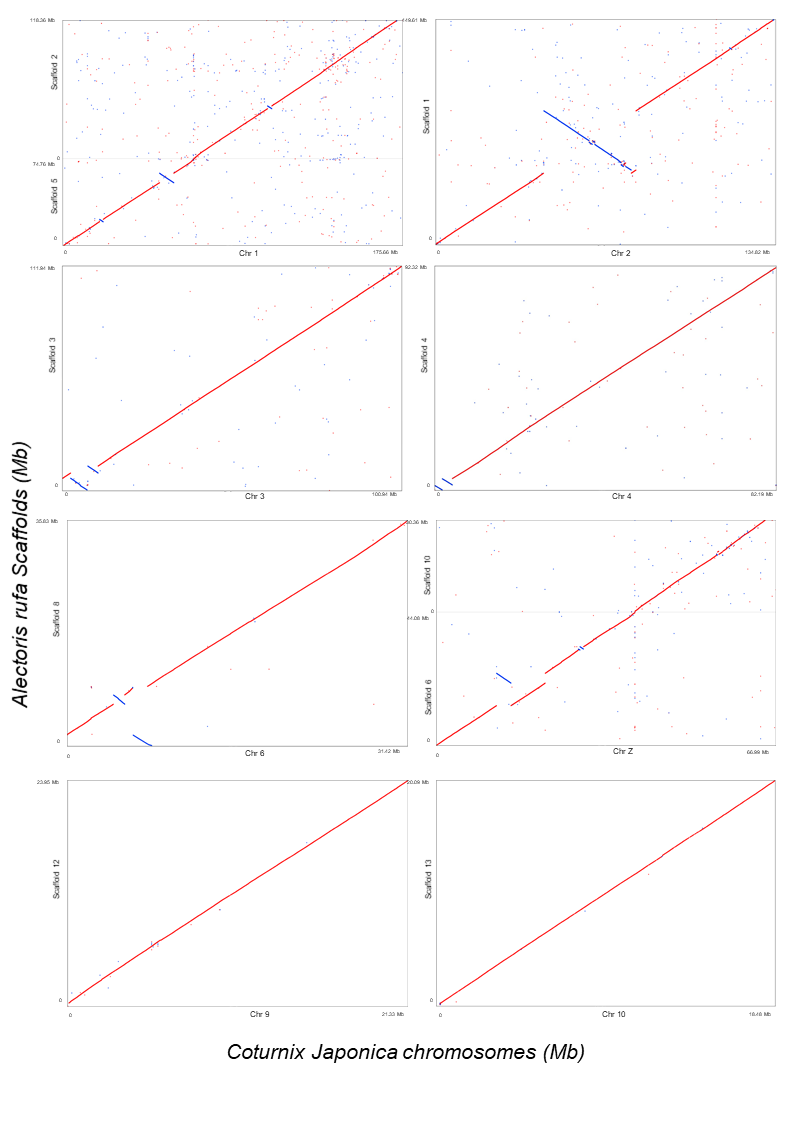

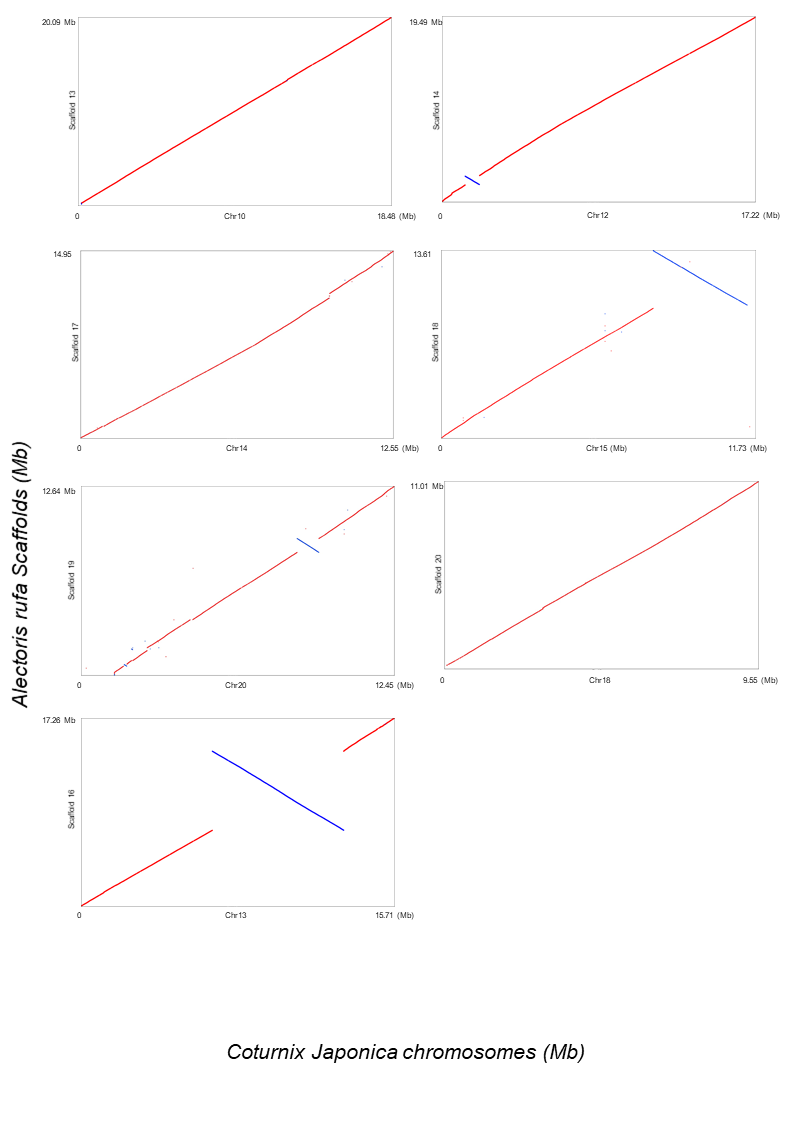

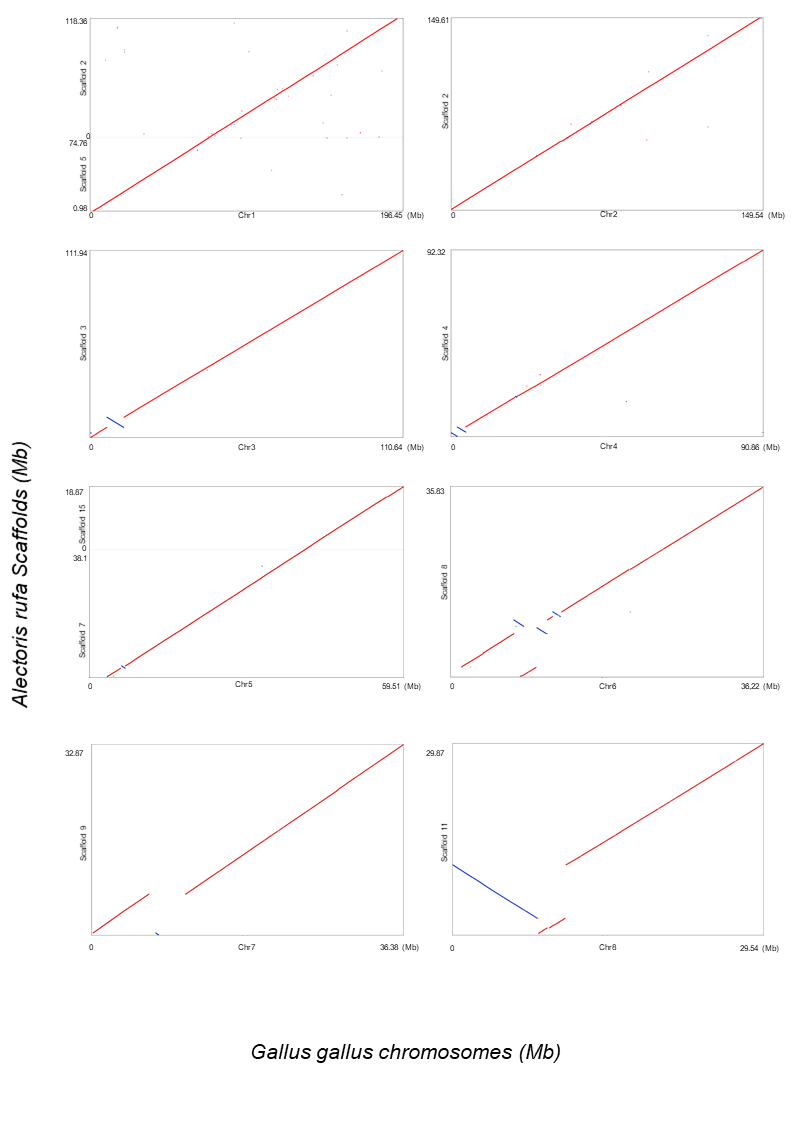

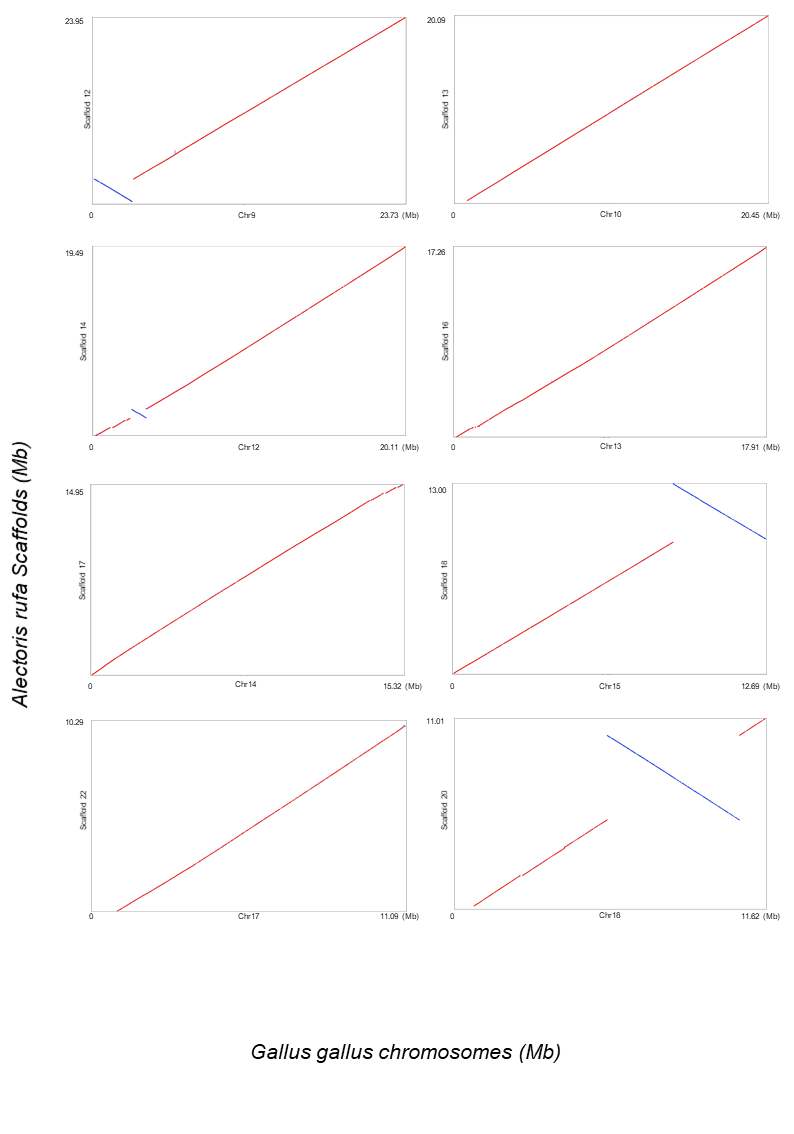

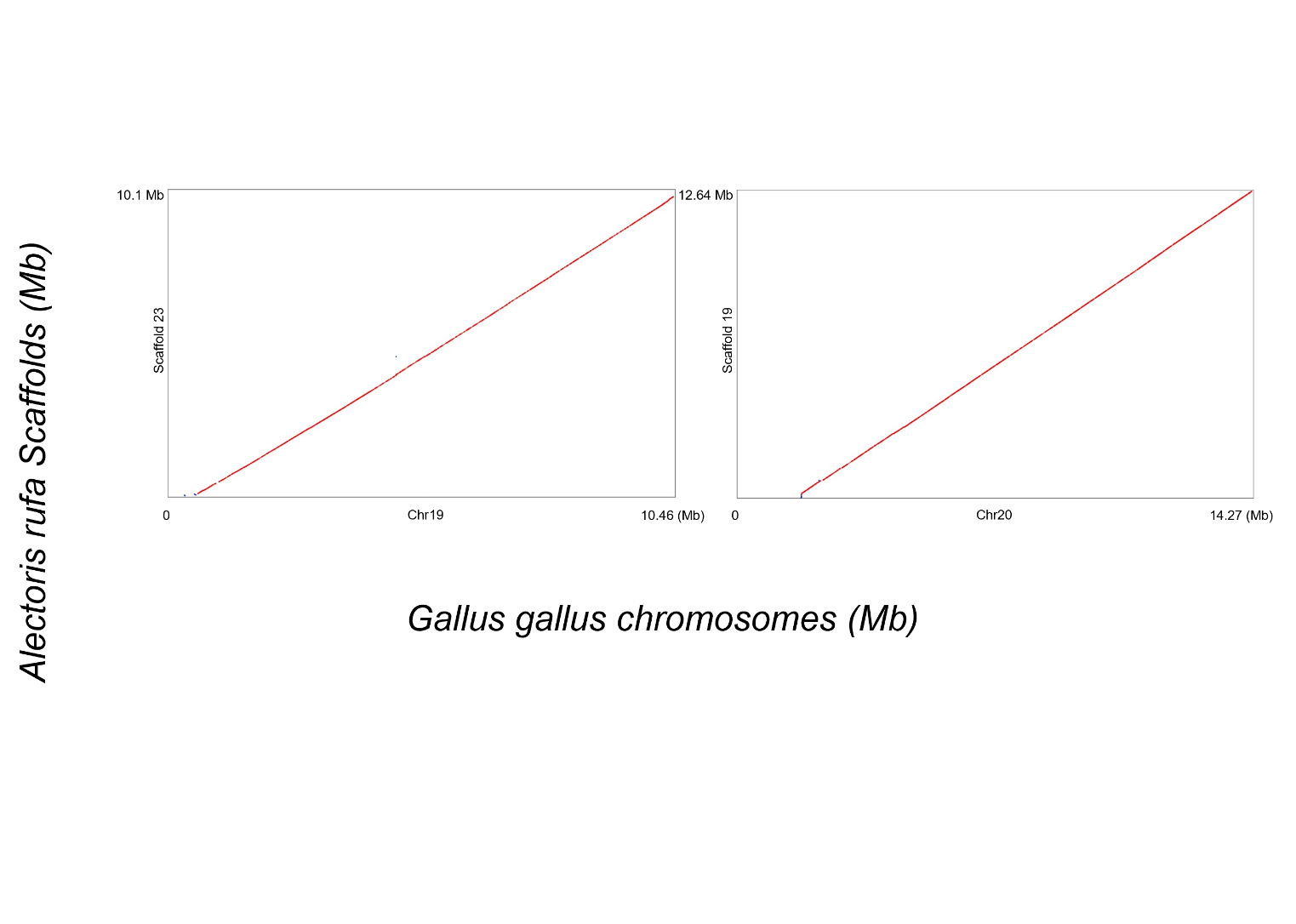

Figure S2: Dot plots showing the alignment of the largest 23 *A. rufa* scaffolds to the reference macro chromosomes of *C. japonica* and *G. gallus*. We represent *A. rufa*’s scaffolds on the y-axis and the chromosomes of *C. japonica* and *G. gallus* on the X-axis. A dot in a plot represents an aligned 10kb block of synteny, with red indicating forward alignment and blue signifying reverse alignment. The axis labels are in Megabase pairs (Mb).

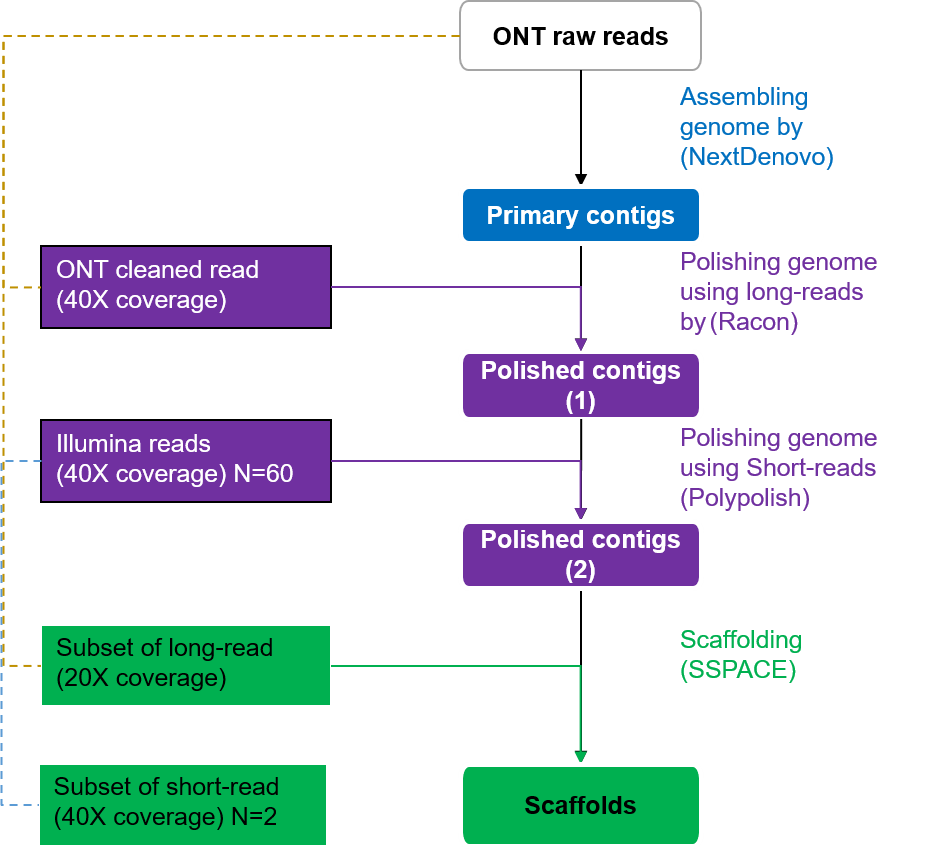

Figure S3: Pipeline for assembling the reference genome of the red-legged partridge (*A. rufa*). Color-coding indicates assembly stages: blue represents primary contig assembly level, purple denotes the genome polishing stage using both long and short-read data, and green highlights the scaffolding process of the polished contig-level assembly with a combination of long and short-reads. N: number of sequenced *A. rufa* individuals.
